# Cell-free Tumor Methylome Analysis of Small Cell Lung Cancer Patients Identifies Subgroups with Prognostic Associations

**DOI:** 10.1101/2022.05.12.491321

**Authors:** Sami Ul Haq, Sabine Schmid, Mansi K. Aparnathi, Katrina Hueniken, Luna Jia Zhan, Danielle Sacdalan, Janice J.N. Li, Nicholas Meti, Devalben Patel, Dangxiao Cheng, Vivek Philip, Ming S. Tsao, Michael Cabanero, Daniel de Carvalho, Geoffrey Liu, Scott V. Bratman, Benjamin H. Lok

## Abstract

**Introduction:** Small cell lung cancer (SCLC) is a highly aggressive type of cancer with a high risk of recurrence. The SCLC methylome may yield biologic insight but is understudied due to difficulty in acquiring primary patient tissue. Here, we comprehensively profile the SCLC methylome using cell-free methylated DNA immunoprecipitation sequencing (cfMeDIP-seq).

**Methods:** cfDNA was extracted from plasma samples collected from 74 SCLC patients prior to initiation of first-line treatment and from 20 non-cancer smoker participants. Genomic DNA (gDNA) was also extracted from paired peripheral blood leukocytes from the 74 SCLC patients and 7 accompanying circulating-tumour-cell patient-derived xenografts (CDX). cfDNA and gDNA were used as input for cfMeDIP-seq. We developed PeRIpheral blood leukocyte MEthylation (PRIME) subtraction as an algorithm to improve tumour specificity of cell-free methylome.

**Results:** SCLC total plasma cfDNA methylation profiles obtained using cfMeDIP-seq are representative of CDX tumour methylation. SCLC cfDNA methylation is distinct from non-cancer plasma. Using PRIME and k-means consensus clustering, we identified two SCLC methylome clusters with prognostic associations. These clusters had methylated biological pathways related to axon guidance, neuroactive ligand−receptor interaction, pluripotency of stem cells, and were differentially methylated at long noncoding RNA, LINEs, SINEs, retrotransposons, and other repeats features.

**Conclusions:** We have comprehensively profiled the SCLC methylome using cfMeDIP-seq in a large patient cohort and identified methylome clusters with prognostic associations. Our work demonstrates the potential of liquid biopsies in examining SCLC biology encoded in the methylome.

## Introduction

Small cell lung cancer (SCLC) is a highly aggressive subset of lung cancer^1^. While often sensitive to first-line therapy, most SCLC patients develop recurrent disease accompanied by therapeutic resistance^1^. Epigenetic mechanisms, specifically DNA methylation, may contribute towards SCLC oncogenesis, recurrence and resistance^1^, through tumourigenesis and epigenetic reprogramming^2^.

Two key SCLC DNA methylome studies examined patient tissue to identify a methylation-defined differentiation block^3^ and unique methylation-defined subtypes^4^. However, progress in comprehensive methylome profiling of SCLC is impeded by lack of primary tumour tissue. Liquid biopsy using cell-free DNA (cfDNA) presents a solution. Previous cfDNA analysis in SCLC evaluated genetic changes, demonstrating a high tumour mutation burden and an increased mutant allele fraction in plasma cfDNA^5^. These findings demonstrate the potential of liquid biopsy analyzes of SCLC tumour biology.

Here, we comprehensively profiled the methylome of plasma cfDNA from SCLC patients to identify novel putative biomarkers and epigenetic mechanisms of disease. For this, we conducted cell-free methylated DNA immunoprecipitation and high-throughput sequencing (cfMeDIP-seq)^6^, which has previously been applied to pancreatic^6^, non-SCLC^6^, renal cell^7^, glioma^8^, head and neck^9^, and other cancers. We analyzed 74 SCLC patients’ plasma samples, enriched for tumour-derived signal with PeRIpheral blood leukocyte (PBL) MEthylation (PRIME) subtraction using matched PBLs, and evaluated biological, clinical, and prognostic associations. Our findings demonstrate the utility of liquid biopsies to examine SCLC methylome and identified two methylation-defined clusters of patients with prognostic association.

## Materials and Methods

cfDNA was extracted from approximately 2mL of plasma using the QIAamp Circulating Nucleic Acid Kit. PBL genomic DNA (gDNA) was extracted using the Roche DNA Isolation Kit for Mammalian Blood followed by sonication and size-selection using Beckman Coulter AMPure XP beads. Patient circulating-tumour-cell-derived xenograft (CDX) models had gDNA extracted from tumour tissue using the Qiagen DNeasy Blood & Tissue Kit. cfMeDIP-seq was done as described previously^10^. 10ng of cfDNA or gDNA was combined with 90ng of lambda filler DNA. CDX models derived as previously described^11^. Methylome data was analyzed by dividing the chromosome 1-22 into 300bp windows. PRIME subtraction was applied to cfDNA methylome as follows: 1) PBL MeDIP data was converted to beta-value using R package MeDEStrand, 2) Median beta value per window calculated for all PBLs, 3) Windows with beta < 0.3 and #CGs per window >= 5 were identified, 4) Plasma cfMeDIP data was subset to these windows. We applied the R packages DESeq2 (differential methylation analysis), ConsensusClusterPlus (consensus clustering), and clusterprofiler (KEGG pathway analysis). Supplemental Data has detailed methods.

## Results

### SCLC cfDNA methylome reflects tumour tissue methylation and is distinct from NCC

We examined the cell-free methylome of 74 SCLC patients using blood samples collected prior to first-line treatment (Figure 1A). 88% of patients (n=66) were current or former smokers and 57% had extensive-stage (ES) SCLC (n=42; Figure 1A) with the remainder limited-stage (LS). We examined the cell-free methylomes of 20 non-cancer control (NCC) participants. NCC participants were smokers and were similar in age and sex to the SCLC cohort (Figure 1A). Detailed demographic characteristics can be found in Supplemental Data Table 1.

**Figure 1.**
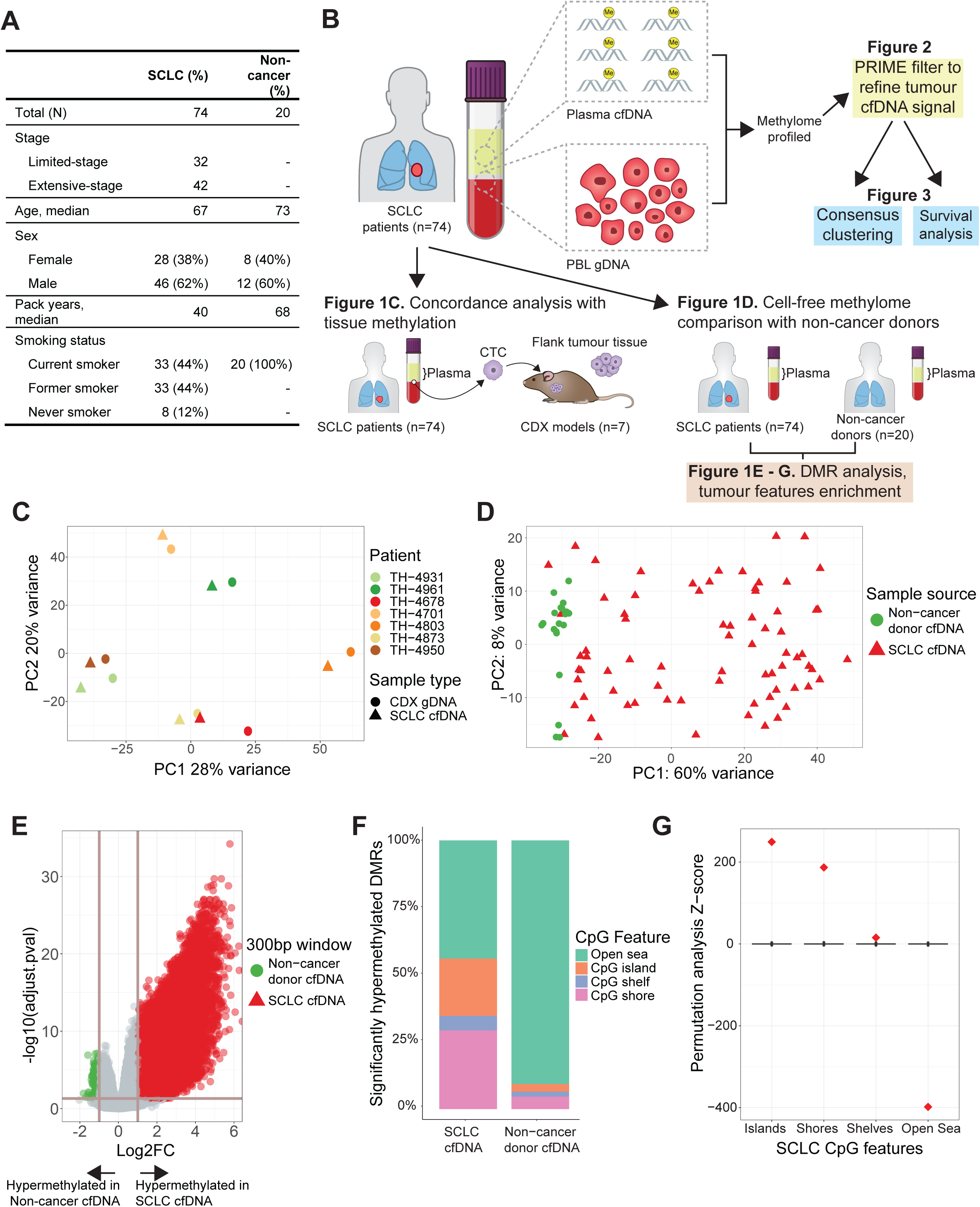
SCLC cell-free methylome reflects tissue and is distinct from non-cancer. A. Demographics and clinical characteristics of 74 small cell lung cancer (SCLC) patients and non-cancer donors. B. Schematic of overall structure of study and major analyses done. C. Principal component analysis of genome-wide methylation profiles of total methylated cell-free DNA (cfDNA) from SCLC patients (n=7) and methylated genomic DNA from paired circulating tumour cell-derived xenograft (CDX) models (n=7). D. Principal component analysis of genome-wide methylation profiles of total methylation cfDNA from SCLC patients (n=74) and non-cancer donors (n=20). E. Volcano plot of differentially methylated region (DMR) analysis of SCLC patients (n=74) and non-cancer donors (n=20). Each dot corresponds to a 300bp region of the genome. The horizontal line corresponds to p-adjusted = 0.05 and vertical lines correspond to log2 fold-change of +/-2. There are 51,666 hypermethylated DMRs in SCLC and 1,019 in non-cancer donors. F. Bar plot of significantly hypermethylated DMRs corresponding to CpG features observed in SCLC patients (n=51,666 DMRs) and non-cancer donors (n=1,019 DMRs). G. Permutation analysis of the hypermethylated DMRs observed in SCLC (n=51,666 DMRs).

To assess whether cfMeDIP-seq can ascertain SCLC tumour tissue methylation, we performed a genome-wide concordance analysis of methylome profiles (n=8.5e6 300bp windows) of cfDNA of SCLC patients (n=7) and gDNA of their respective CDX tumour tissue (n=7). Patient cfDNA was highly concordant with CDX methylation by principal component analysis (PCA; Figure 1C) and strongly correlated by correlation analysis of normalized read counts within each window (median r=0.92, n=7). Thus, cfMeDIP-seq data appeared representative of tissue-level DNA methylation.

However, as others have reported SCLC patients that successfully engraft CDXs have increased CTCs which could increase tumor cfDNA^12^. For our SCLC patients without a corresponding CDX (n=67), the contribution of non-cancer cfDNA is expected to be higher. Therefore, we assessed if cfMeDIP-seq could distinguish between SCLC and NCC methylation through examining genome-wide cfDNA methylome profiles by PCA (Figure 1D). NCCs were distinct from SCLC suggesting cfMeDIP-seq distinguished non-cancer methylation from cancer. Through differentially methylation region (DMR) analysis between SCLC and NCC, 51,666 and 1,019 significantly hypermethylated DMRs were identified in SCLC and NCC, respectively (p-adj < 0.05; log2FC >1; Figure 1E). SCLC significant DMRs were enriched in CpG islands and shores relative to NCCs, whereas NCC significant DMRs were enriched in open-sea regions (Figure 1F). Permutation analysis revealed that CpG features were significantly enriched in SCLC cfDNA DMRs and verified that cfMeDIP-seq is tumour-specific (Figure 1G).

### PRIME removes non-cancer methylation using paired PBLs from same SCLC patients

Non-tumour methylation signal in the SCLC cfDNA data was examined and quantified using MethylCIBERSORT. This allowed us to approximate the proportion of methylation in plasma cfDNA from non-tumour cells. We found PBLs to be a large contributor to plasma cfDNA methylation (Supplemental Data Fig. 1-5).

To increase specificity of the SCLC signal to non-cancer noise ratio, we implemented a novel approach utilizing paired PBL gDNA collected from the same SCLC patient plasma source material at identical timepoints (Figure 2A). Comparison of total plasma cfDNA to PBL gDNA by PCA, revealed that PBLs exhibited a distinct methylation signal (Figure 2B). Next, we examined the methylome of SCLC total plasma cfDNA alone (Figure 2C), and after applying our novel algorithmic filter, PRIME, to reduce PBL methylation signals (Figure 2D). PRIME filtered out non-tumour noise in the cfDNA, reducing 8.5e6 windows to 1.9e5 SCLC-specific windows. By PCA, PRIME increased the variance explained by methylation in PC1 from 40% (Figure 2C) to 57% (Figure 2D) and decreased the variance in PC2 from 7% (Figure 2C) to 4% (Figure 2D). PRIME also identified two distinct PCA groups (Figure 2D), refined our cfDNA methylation, and increased cfMeDIP-seq SCLC-specificity.

**Figure 2.**
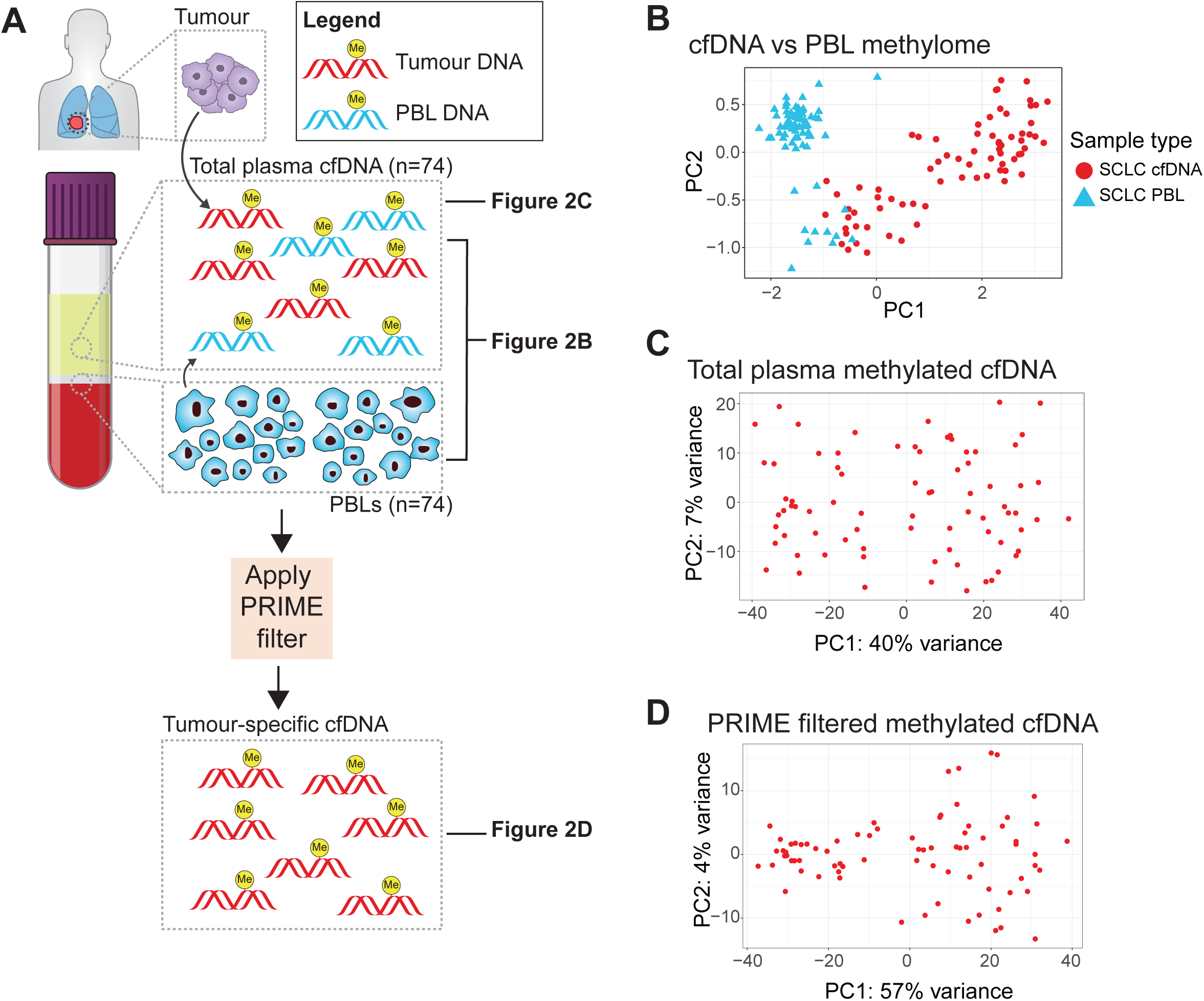
Filtering out non-cancer methylation signal from plasma cfDNA. A. Overall schematic outlining our approach to first compare SCLC total cfDNA and SCLC PBL gDNA methylome (2B), SCLC total cfDNA methylome (2C), and PRIME-filtered cfDNA methylome (2D). B. Principal component analysis (PCA) of methylation profiles of 74 SCLC patients comparing total plasma cfDNA methylation to PBL gDNA methylation. For each patient, plasma cfDNA and PBL gDNA are extracted using peripheral blood samples collected prior to starting first-line chemotherapy. Genome-wide methylation profiles are examined in this PCA plot (∼8.5e6 300bp windows). Red filled circles correspond to cfDNA samples and blue triangles are PBL samples. C. PCA of total plasma cfDNA methylation profiles of the 74 SCLC patients. Genome-wide methylation profiles are examined in this PCA plot (∼8.5e6 300bp windows). Red filled circles correspond to cfDNA samples. D. PCA of PRIME filtered plasma cfDNA methylation profiles of the 74 SCLC patients. In this PCA, methylation profiles for ∼190,000 300bp windows are examined.

### Identifying methylation-defined prognostic clusters and examining differentially methylated pathways and features

To define potential SCLC methylome subgroups, we applied k-means consensus clustering on the PRIME-filtered cfDNA methylome profiles for all 74 SCLC patients. We determined 2 methylation-defined clusters was the optimal number (Supplemental Data Fig 6), which we designated as Clusters A and B (Figure 3A). Patients in these Clusters had significantly different overall survival (OS; HR = 2.02, p = 0.014; Figure 3B) where Cluster B had a median OS of 13 months compared to 21 in Cluster A. However, after adjusting for stage, Clusters A and B were not significant (Supplemental Data Table 2). Interestingly, after we stratified stage by Cluster, Cluster B consistently had worse OS in either stage, which was more apparent in LS-SCLC (Supplemental Data Fig 7). Since all blood samples were collected prior to treatment, these methylation-defined clusters may hold prognostic potential (Figure 3B) and methylome associated biologic consequence.

**Figure 3.**
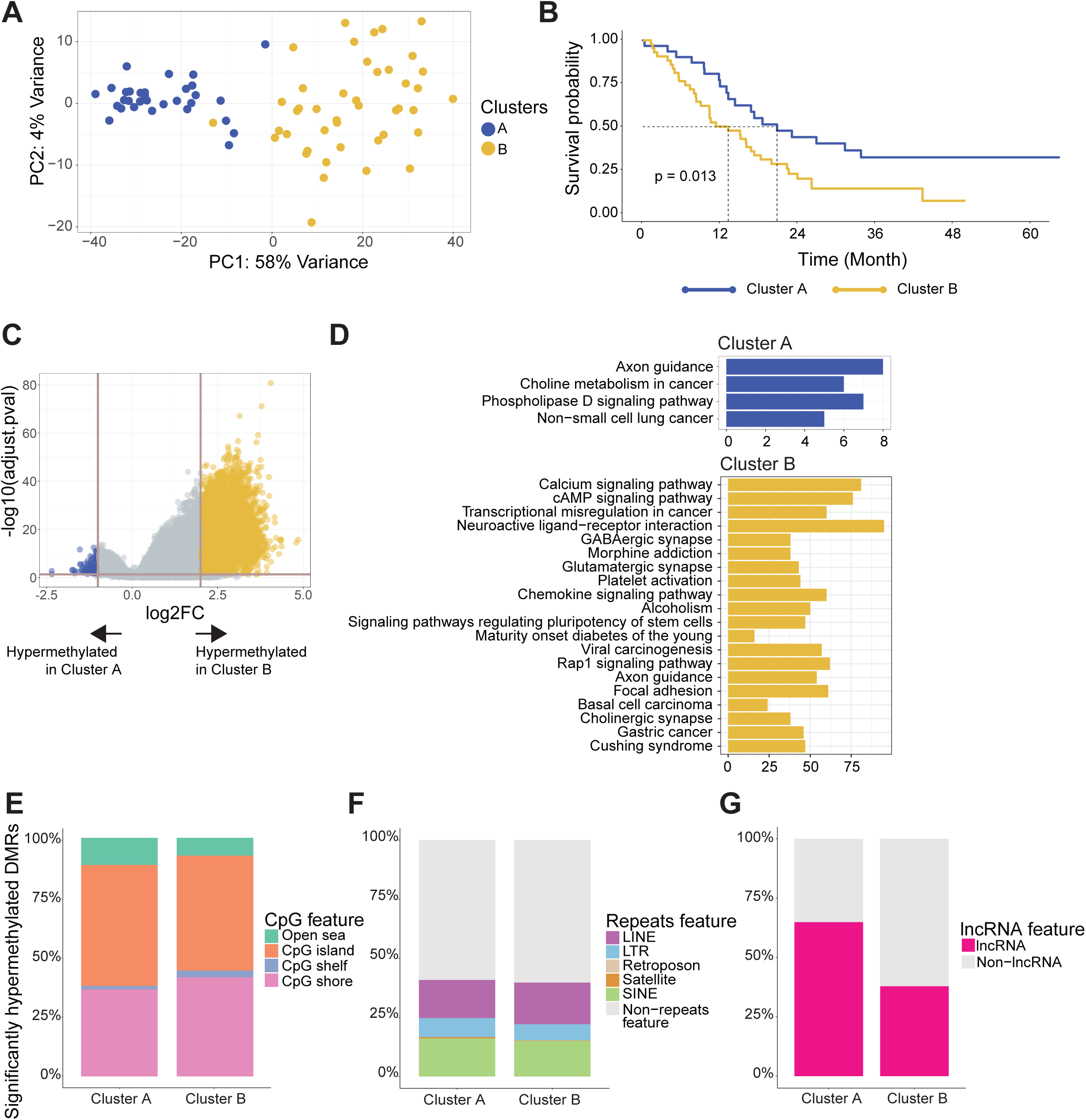
Identifying prognostic clusters and examining differentially methylated pathways and features. A. Consensus clustering done on PRIME-filtered SCLC methylated cfDNA identified two clusters, A and B. B. Kaplan-Meier survival analysis on cluster A and B identified by consensus clustering. C. Volcano plot of differentially methylated region (DMR) analysis between cluster A and B. The horizontal line corresponds to p-adjusted = 0.05 and vertical lines correspond to log2 fold-change of +2 and -1. There are 174 hypermethylated DMRs in cluster A and 9037 in cluster B. D. KEGG pathway analysis of hypermethylated DMRs in cluster A and cluster B. E. Bar plots of significantly hypermethylated DMRs observed in Cluster A (n=9037) and Cluster B (n=174) corresponding to CpG features. F. Bar plots of significantly hypermethylated DMRs observed in Cluster A (n=9037) and Cluster B (n=174) corresponding to repeats features. G. Bar plots of significantly hypermethylated DMRs observed in Cluster A (n=9037) and Cluster B (n=174) corresponding to long-noncoding RNA (lncRNA).

Subsequently, we conducted a DMR analysis on Clusters A and B (Figure 3C). This analysis identified 174 significantly hypermethylated DMRs in Cluster A (p-adj < 0.5 log2FC <-1) and 9,037 in Cluster B (p-adj < 0.05 log2FC > 2). Cluster A and Cluster B had similar hypermethylated DMR proportions among all CpG features (Figure 3E). Interestingly, approximately 40% of all significantly hypermethylated DMRs in both Cluster A and B also corresponded to features such as short interspersed nuclear elements (SINEs), long interspersed nuclear elements (LINEs), long terminal repeats (LTRs), retrotransposons, and satellites (Figure 3F). 65% percent of significantly hypermethylated DMRs in Cluster A corresponded to long non-coding RNA (lncRNA) windows compared to 38% in Cluster B (Figure 3G). This suggested a possible role of lncRNAs in mediating the prognostic associations identified in Cluster A and B.

To understand methylation differences in terms of biological pathways, a KEGG pathway analysis was performed on the significantly hypermethylated DMRs (Figure 3D). In Cluster A, 174 significant DMRs corresponded to 137 genes whereas in Cluster B, 9,037 DMRs corresponded to 2,131 genes. Pathways corresponding to axon guidance or phospholipase D signaling pathway, or non-small cell lung cancer, were enriched in Cluster A (Figure 3D). On the other hand, Cluster B had several pathways like neuroactive ligand−receptor interaction, immune chemokine signaling, and pathways regulating pluripotency of stem cells (Figure 3D).

## Discussion

Our study demonstrated the utility of the cfMeDIP-seq assay in studying the methylome of SCLC and demonstrated that plasma methylation is representative of tissue (Figure 1C) and is enriched in regions that are cancer specific (Figure 1D, Figure 1E)^13^. To our knowledge, this is the largest study to comprehensively examine the DNA methylation of SCLC patients by whole genome analysis. In contrast to prior studies^3,4^, our genome-wide approach profiled methylated DNA loci beyond 450K/EPIC array probes and identified hypermethylated non-coding and repeat elements (e.g. LINE, SINE, lncRNA, etc.). In addition, our available clinical annotation with these treatment naïve patient blood samples allowed us to correlate tumour methylation with clinical outcomes.

Previous liquid biopsy studies have observed PBL being major contributors to the cfDNA signal^14,15^. In our data, we quantified this non-cancer signal using MethylCIBERSORT underscoring the need to filter the non-cancer contribution to plasma cfDNA. Currently, most liquid biopsy studies do not control for PBL cfDNA and this may affect the tumour-specificity of the resultant methylome analysis and impact certain application goals. Using PRIME, a bespoke PBL algorithmic to refine SCLC-specific methylome signal using paired PBLs from the same patients, we increased tumour-specificity of our resultant cfMeDIP-seq data (Figure 2D). We identified two Clusters, A (better OS) and B (worse OS). This prognostic association suggests that methylation may be contributing to cancer progression and metastasis – highlighting the need to unravel SCLC biology that is mediated by epigenetic mechanisms.

DMR analysis of these Clusters identified repeat features such as LINEs/SINEs and lncRNA not previously comprehensively characterized in the SCLC methylome. Methylation of these regions may implicate them in epigenetically mediating biological pathways in SCLC and may elucidate novel biologic insights worthy of future investigation.

cfMeDIP-seq allows interrogation of the SCLC methylome in a non-invasive, comprehensive manner. cfMeDIP-seq can be applied longitudinally to interrogate the SCLC methylome using pre-, on-, post-treatment samples at a scale and accessibility not possible by invasive tissue collection methods. Ultimately, cfMeDIP-seq can be used to identify treatment-induced changes in SCLC that may be contributing to resistance. By unraveling the biology of these potential epigenetic mechanisms, we hope to continually improve the outcomes of our patients with SCLC.

## Supporting information

Supplemental Data - Detailed KEGG Pathway Analysis

## Works Cited

1. Rudin CM, Brambilla E, Faivre-Finn C, Sage J. Small-cell lung cancer. Nat Rev Dis Prim. 2021;7(1):3. doi:10.1038/s41572-020-00235-0

2. Kulis M, Esteller M. DNA methylation and cancer. Adv Genet. 2010;70:27–56. doi:10.1016/B978-0-12-380866-0.60002-2

3. Kalari S, Jung M, Kernstine KH, Takahashi T, Pfeifer GP. The DNA methylation landscape of small cell lung cancer suggests a differentiation defect of neuroendocrine cells. Oncogene. 2013;32(30):3559–3568. doi:10.1038/onc.2012.362

4. Poirier JT, Gardner EE, Connis N, et al. DNA methylation in small cell lung cancer defines distinct disease subtypes and correlates with high expression of EZH2. Oncogene. 2015;34(48):5869–5878. doi:10.1038/onc.2015.38

5. Almodovar K, Iams WT, Meador CB, et al. Longitudinal Cell-Free DNA Analysis in Patients with Small Cell Lung Cancer Reveals Dynamic Insights into Treatment Efficacy and Disease Relapse. J Thorac Oncol. 2018;13(1):112–123. doi:10.1016/j.jtho.2017.09.1951

6. Shen SY, Singhania R, Fehringer G, et al. Sensitive tumour detection and classification using plasma cell-free DNA methylomes. Nature. 2018;563(7732):579–583. doi:10.1038/s41586-018-0703-0

7. Nuzzo PV, Berchuck JE, Korthauer K, et al. Detection of renal cell carcinoma using plasma and urine cell-free DNA methylomes. Nat Med. 2020;26(7):1041–1043. doi:10.1038/s41591-020-0933-1

8. Nassiri F, Chakravarthy A, Feng S, et al. Detection and discrimination of intracranial tumors using plasma cell-free DNA methylomes. Nat Med. 2020;26(7):1044–1047. doi:10.1038/s41591-020-0932-2

9. Burgener JM, Zou J, Zhao Z, et al. Tumor-Naïve Multimodal Profiling of Circulating Tumor DNA in Head and Neck Squamous Cell Carcinoma. Clin Cancer Res. 2021;27(15):4230–4244. doi:10.1158/1078-0432.CCR-21-0110

10. Shen SY, Burgener JM, Bratman S V, De Carvalho DD. Preparation of cfMeDIP-seq libraries for methylome profiling of plasma cell-free DNA. Nat Protoc. 2019;14(10):2749–2780. doi:10.1038/s41596-019-0202-2

11. Hodgkinson CL, Morrow CJ, Li Y, et al. Tumorigenicity and genetic profiling of circulating tumor cells in small-cell lung cancer. Nat Med. 2014;20(8):897–903. doi:10.1038/nm.3600

12. Vickers AJ, Frese K, Galvin M, et al. Brief report on the clinical characteristics of patients whose samples generate small cell lung cancer circulating tumour cell derived explants. Lung Cancer. 2020;150:216–220. doi:10.1016/j.lungcan.2020.11.002

13. Yates J, Boeva V. Deciphering the etiology and role in oncogenic transformation of the CpG island methylator phenotype: a pan-cancer analysis. Brief Bioinform. 2022;23(2). doi:10.1093/bib/bbab610

14. Croitoru VM, Cazacu IM, Popescu I, et al. Clonal Hematopoiesis and Liquid Biopsy in Gastrointestinal Cancers. Front Med. 2021;8:772166. doi:10.3389/fmed.2021.772166

15. Chan HT, Nagayama S, Chin YM, et al. Clinical significance of clonal hematopoiesis in the interpretation of blood liquid biopsy. Mol Oncol. 2020;14(8):1719–1730. doi:10.1002/1878-0261.12727

