## Supplemental Data - Extended Methods for "Cell-free Tumor Methylome Analysis of Small Cell Lung Cancer Patients Identifies Subgroups with Prognostic Associations"

**cfDNA Extraction**

Peripheral blood collected in EDTA tubes was first spun down at 4000xg at 4 degrees Celsius for 10 minutes. Subsequently, the top plasma layer was transferred to 15ml Falcon tubes and spun again at 16000xg at 4 degrees Celsius for 10 minutes. The supernatant was then transferred to 1.5ml Eppendorf tubes and stored at -80 degrees Celsius. For cfDNA extraction, approximately 3ml of the processed plasma was used with the QIAamp Circulating Nucleic Acid Kit (Qiagen, cat. no. 55114) as described in this protocol (Shen et al. 2018). Concentration of extracted cfDNA was quantified using the Qubit dsDNA HS Assay Kit (Thermo Fisher Scientific, cat. no. Q32851).

**Peripheral Blood Leukocyte (PBL) DNA Extraction**

Peripheral blood collected in EDTA tubes was first spun down at 4000xg at 4 degrees Celsius for 10 minutes. Subsequently, the buffy coat layer was transferred to a 1.5ml Eppendorf and stored at -80 degrees Celsius. Genomic DNA was extracted using the Sigma-Aldrich DNA Isolation Kit for Mammalian Blood (Roche, cat. no. 11667327001) and then sonicated to 150bp using the Covaris M220 Focused-ultrasonicator. Sonicated DNA was then size selected using Agencourt AMPure XP beads (Beckman Coulter, cat. no. A63881) using a 0.8x ratio.

**CDX Generation**

Circulating tumour cells (CTCs) were extracted from one EDTA tube of peripheral blood. The blood was first incubated with 50ul RosetteSep^TM^ (cat. no. 15705) per ml of patient blood for 20 minutes in a rotator at room temperature. After 20 min, blood was diluted with equal volume of wash buffer (WB; 10% HITES media in HBSS) and layered on top of 15 ml of Ficol-Plaque Plus in a 50 ml SepMate^TM^ tube (cat. no. 85450), followed by centrifugation at 1200g for 10 minutes. Contents above plastic insert of SepMate^TM^ tube were collected in fresh 50ml tube, 30 ml of WB was added, followed by another centrifugation at 300g for 10 minutes. The supernatant was discarded, and the pellet was resuspended in 3 ml of 1x StemCell Technologies' RBC lysis buffer (cat. no. 20120) and incubated for 10 minutes at room temperature. Subsequently, 10 ml of WB was added, followed by centrifugation at 300g for 10 minutes. Finally, the supernatant was removed, and the pellet was resuspended in 100ul of 50% of Matrigel (cat. no. 354230) in HITES media. CTCs in HITES/Matrigel mixture were injected subcutaneously into flank of 6 –16-week immunocompromised (NSG) mice.

**CDX gDNA extraction**

CDX tumour tissue gDNA was extracted using the DNeasy Blood & Tissue Kit (Qiagen, cat. no. 69504). Up to 25mg of tumour tissue was used. The extracted gDNA was then sonicated and size-selected in an identical manner as described in the “Peripheral Blood Leukocyte (PBL) DNA Extraction” section.

**cfMeDIP Library Preparation**

cfMeDIP libraries were made using either extracted cfDNA or processed genomic DNA according to the following . For all samples, 10ng sample DNA as input and 90ng of methylated/unmethylated lambda filler DNA was used.

**Next-generation Sequencing**

cfMeDIP-seq libraries were sequenced on an Illumina NovaSeq 6000 instrument (2 × 100 bp paired-end reads) according to the manufacturer’s recommendations using NovaSeq 6000 SP Reagent Kit v1.5. (Illumina, San Diego, CA, USA). All cfMeDIP-seq libraries were sequenced to 100 million reads.

**Processing Sequenced Reads**

Fastq files were processed as follows. First, terminal adaptor sequences were removed using TrimGalore! (version 0.6.5) and aligned to the reference human genome (hg19) using Burrows-Wheeler Alignment tool (BWA; version 0.7.17). Resulting SAM files were then indexed, and duplicate reads removed using SAMtools (version 1.12).

**PRIME Filter**

For all PBL MeDIP libraries, sequenced reads were binned into 300bp windows spanning the entire human genome, excluding ENCODE blacklisted regions. For each 300bp window, beta-value was estimated using the R package MeDEStrand (version 0.0.0.9000). PBL windows with a median beta-value of less than 0.3 (across all PBL samples) and CG density per window greater than or equal to five were selected. These windows (n=1.9e5) are referred to as PRIME-filtered windows.

**Differentially Hyper-Methylated Regions (DMR) analysis**

DMR analysis was done using the R package DESeq2 (version #1.30.1) (Love, Huber, and Anders 2014). For any DMR analysis, samples of interest were divided up into two groups. Sequenced reads were binned into 300bp windows covering the entire human genome. For each window, a normalization factor was applied where read counts for each sample was divided by the mean read count of all samples (Normalization Factor = sample count / Mean of all samples). Bayesian statistical approaches were used to minimize within sample variation and bring extreme values close to the mean, per window. Subsequently, for each window, a general linear model (approximated as y = bx + error) was fitted using the negative binomial distribution (Love, Huber, and Anders 2014).

**Principal component analysis (PCA)**

PCA was done using the built-in plotPCA() function of DESeq2 on counts data from processed samples. For whole-genome methylation profile analysis, 300bp windows corresponding to ENCODE blacklist regions were removed and the remaining windows (n=8.5e6) were examined by PCA. For PRIME-filtered methylation profile analysis, counts data was subset to windows corresponding to PRIME-filtered windows (n=1.9e5). Subsequently, counts data was transformed by variance stabilizing transformation to produce transformed data on the log2 scale and this was normalized with respect to library size. The transformed counts data were visualized by PCA using plotPCA().

**Annotation of genomic regions**

The R package annotatr (1.16.0) was used to annotate genomic regions (300bp windows). Specifically, annotatr was used to annotate CpG features, i.e. if a window falls in a CpG island/shelf/shore/open-sea region or gene features (i.e. promoter, 3’/5’ UTR, exon). ENSEMBL release-104 was used to annotate non-coding gene features such as lncRNA. UCSC RepeatMasker (version: 2021-09-03) was used to annotate repeats features such as LINEs, SINEs, LTRs, retrotransposons, and satellites features.

**Consensus clustering**

To identify methylation-defined biologically relevant subtypes of SCLC, consensus clustering was applied to sequenced read data in the following manner. For each sample, read data was subset to PRIME-filtered windows. The median absolute deviation was calculated for each window, and the top 5000 most deviant windows were selected. Consensus clustering was performed on these top 5000 windows. Consensus clustering was done for k values of 2 to 20, with 1000 resamplings.

**Permutation analysis**

To calculate significance of CpG feature enrichment, permutation analysis was done. First, the human genome was segmented into 300bp windows spanning the entire genome (except sex chromosomes). 51,666 windows were randomly sampled (because this is the number of hypermethylated DMRs observed in SCLC) and the occurrence of CpG feature (i.e. CpG island) was calculated. This was repeated 1000 times. Subsequently, the 1000x calculated frequencies were converted into Z-scores and a null distribution was determined. The observed CpG feature frequency was converted into a Z-score and statistical analysis was done.

**KEGG pathway analysis**

The R package clusterprofiler (version 3.16.1) was used to perform pathway analysis on methylated regions. Gene symbols corresponding to significantly hypermethylated 300bp windows were determined using annotatr. Subsequently, clusterprofiler function enrichKEGG() was run on the gene symbols list.

**Patient Clinical/Demographics and Kaplan-Meier Survival Analysis**

The clinico-demographic features were summarized descriptively, using median and IQR and for continuous variables, and counts and percentages for ordinal/categorical variables. P-values were obtained via fisher's exact test for categorical/ordinal variables and Kruskal Wallis H test for continuous variables.

Overall survival (OS) was defined as the time from small-cell lung cancer diagnosis until death from any cause or censored at last follow-up. Kaplan-Meier curves and log-rank tests were used to visualize survival differences between groups. The association between cluster and overall survival stratified by VA stage at diagnosis was explored using Cox proportional-hazard model. Hazard ratios (HRs) were reported with 95% confidence intervals. The proportionality assumption was met via assessing Schoenfeld residuals against time. Statistical analyses were performed using R software (version 4.1.0).
