## Supplemental Data - Figures & Tables for "Cell-free Tumor Methylome Analysis of Small Cell Lung Cancer Patients Identifies Subgroups with Prognostic Associations"

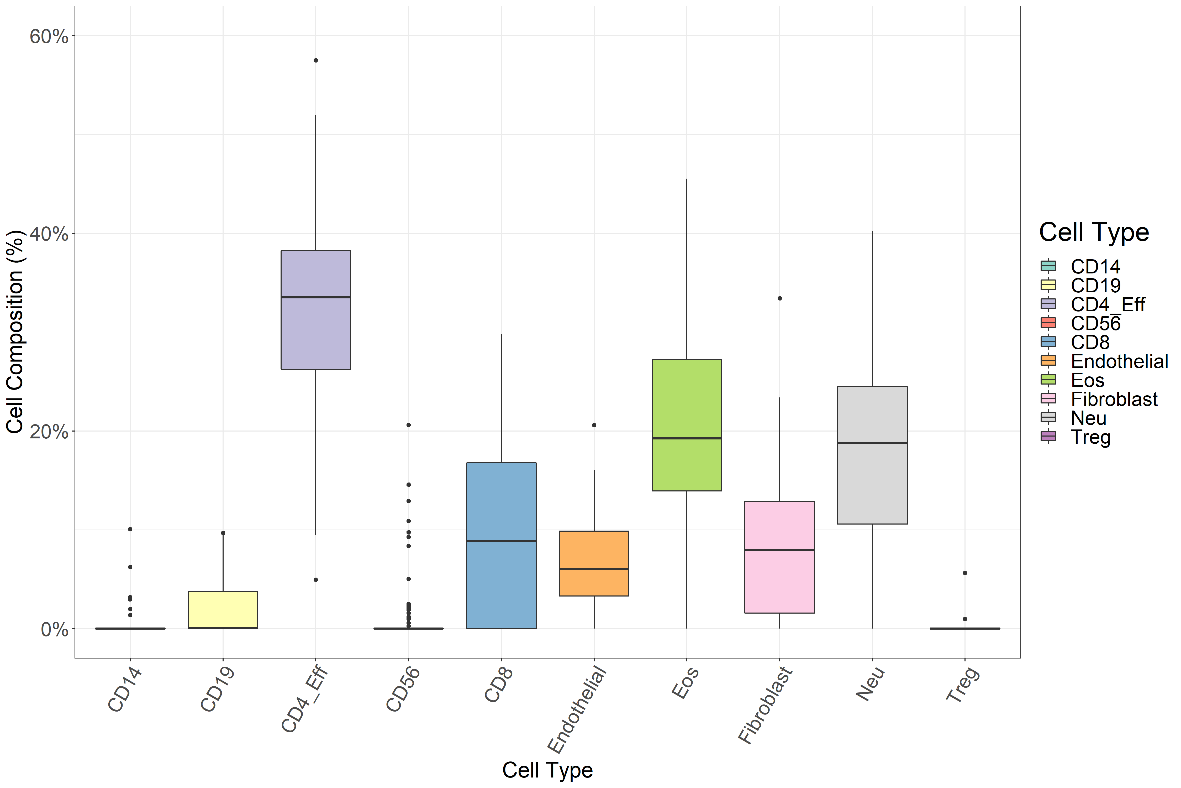

**Supplemental Figure 1.** MethylCIBERSORT composition analysis by total plasma cfDNA methylation, quantifying immune cell presence in the plasma by cell-type. Each box-plot represents the range of cell-composition (%) by cell type for all 74 SCLC patients.

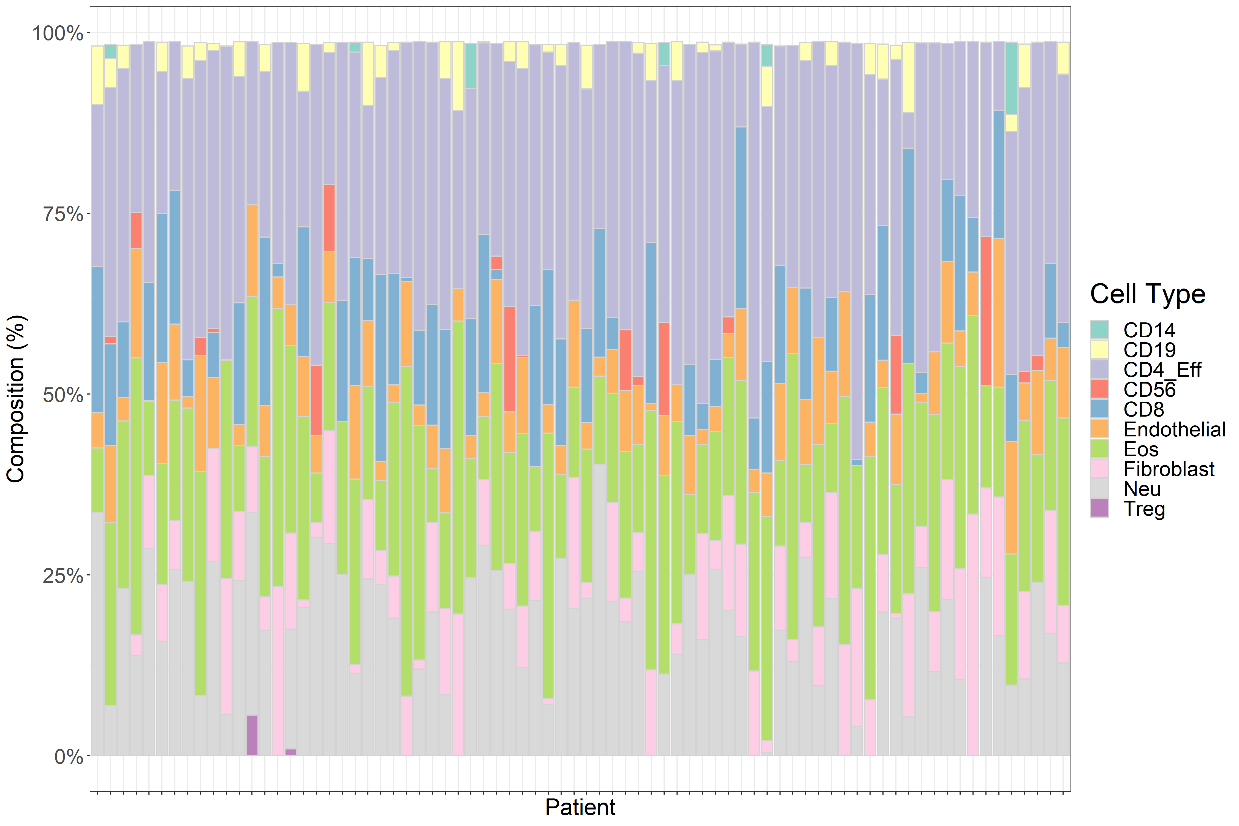

**Supplemental Figure 2.** MethylCIBERSORT composition analysis by total plasma cfDNA methylation, quantifying immune cell presence in the plasma by patient. Each stacked bar-plot represents each of the 74 SCLC patients.

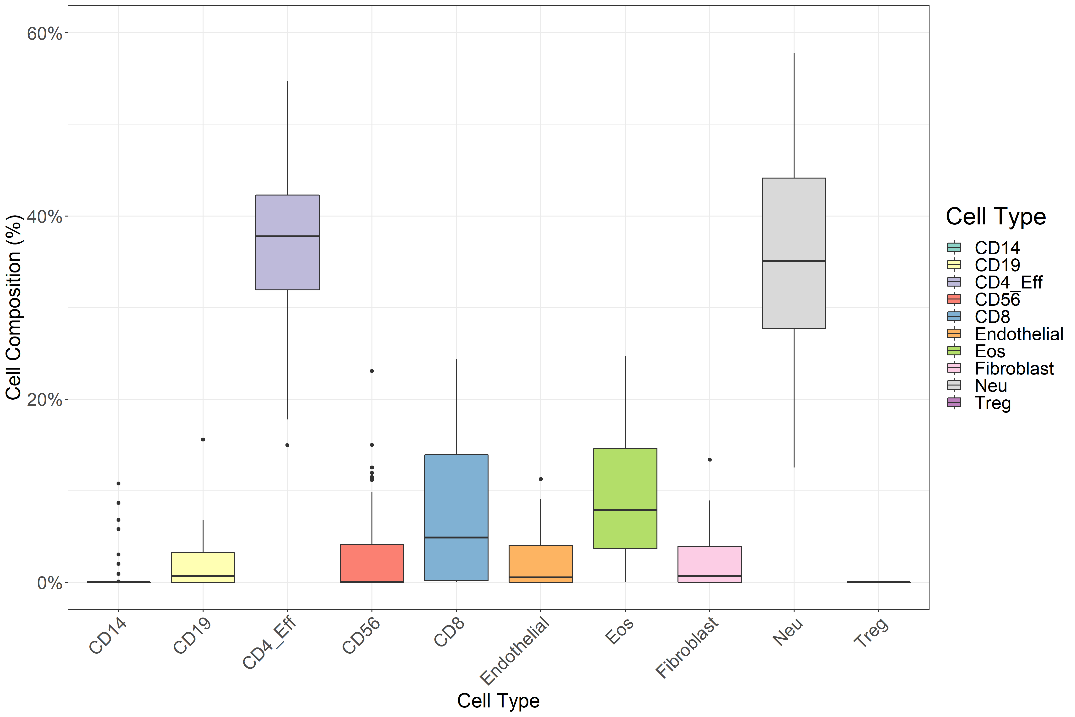

**Supplemental Figure 3.** MethylCIBERSORT composition analysis by PBL gDNA methylation, quantifying immune cell presence in the plasma by cell-type. Each box-plot represents the range of cell-composition (%) by cell type for all 74 SCLC patients.

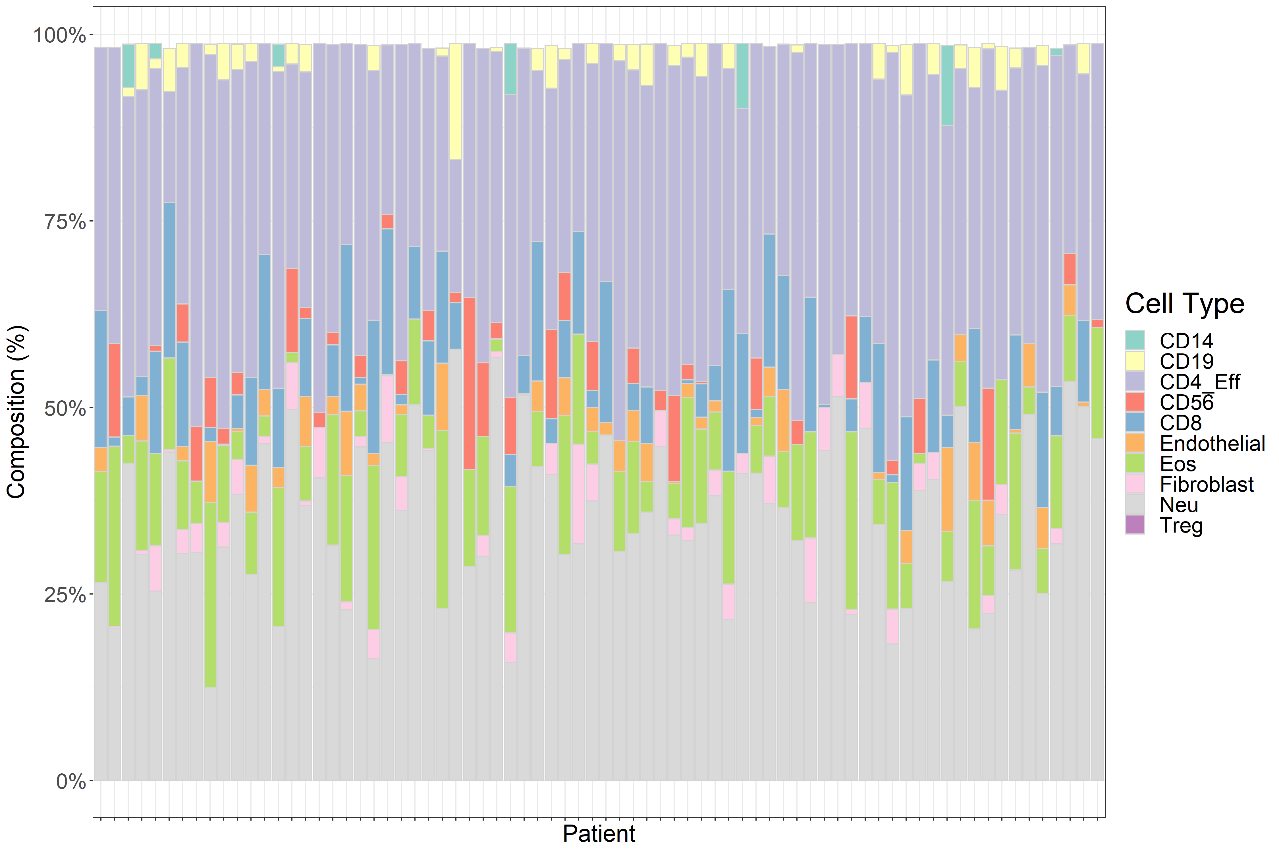

**Supplemental Figure 4.** MethylCIBERSORT composition analysis by PBL gDNA methylation, quantifying immune cell presence in the plasma by individual patient. Each stacked bar-plot represents each of the 74 SCLC patients.

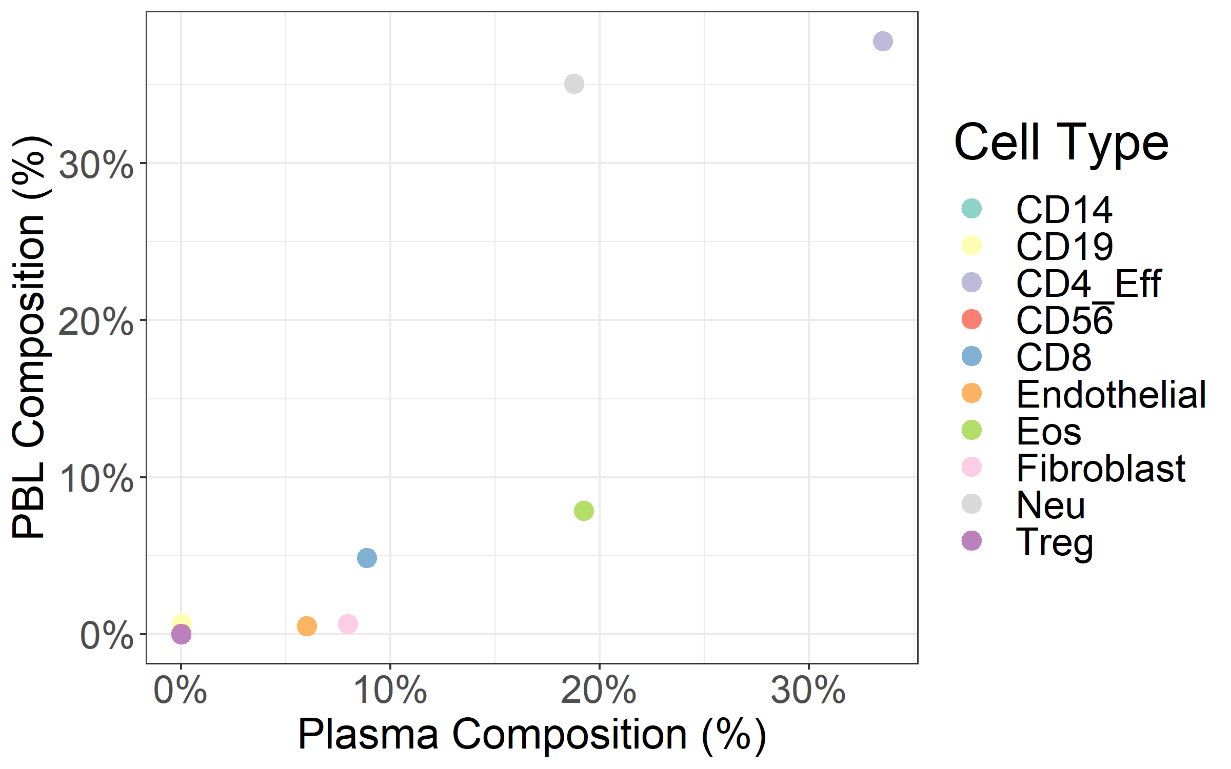

**Supplemental Figure 5.** Examining MethylCIBERSORT quantification of median immune cell signal cell composition (%) by methylation in SCLC total plasma and SCLC PBLs. Each dot represents an immune cell type and the x/y-axis represent the median immune cell composition (%) in either PBL gDNA or total plasma cfDNA for that cell type.

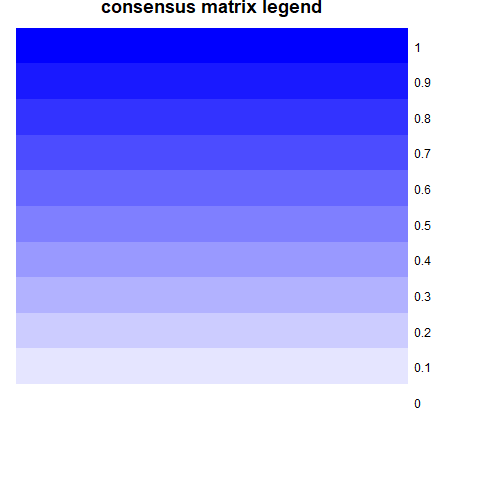

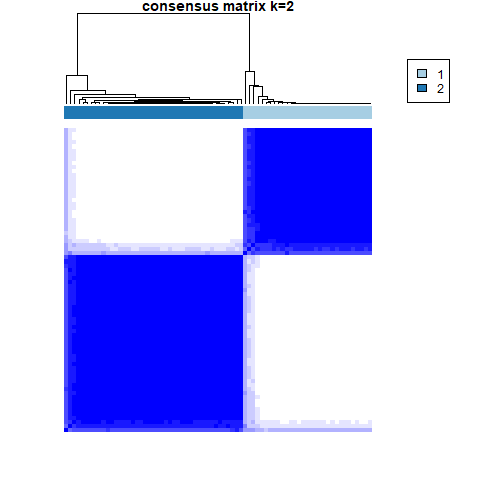

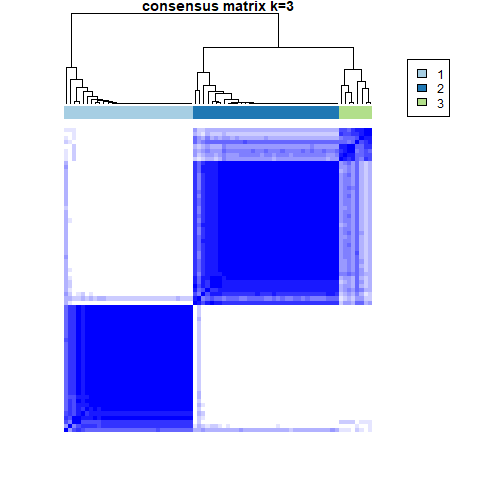

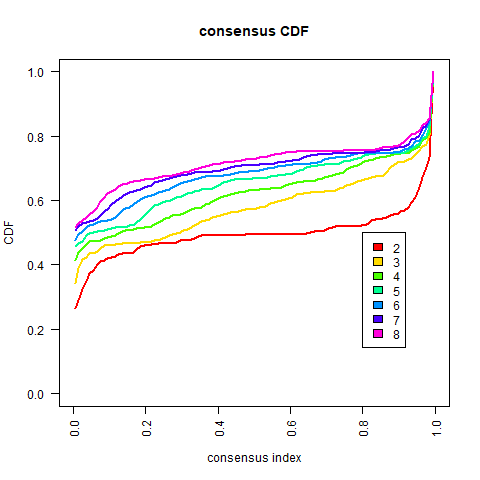

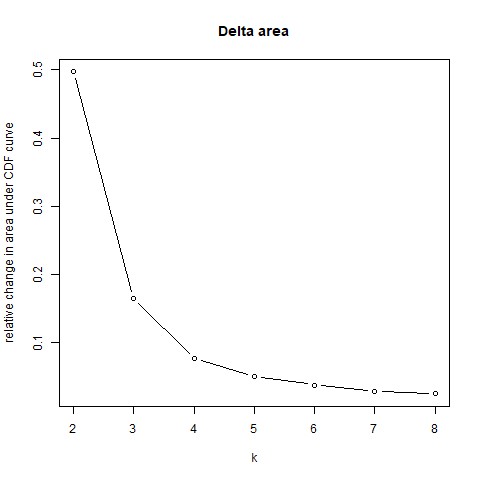

**Supplemental Figure 6**. Consensus clustering done on the PRIME-filtered cfDNA methylome data. Top images compare k=2 vs k=3 consensus clusters. For each k, consensus matrix plots depict consensus values on a white (less consensus) to blue (more consensus) colour scale, are ordered by the consensus clustering, which is shown as a dendrogram, and have samples’ consensus clusters annotated. The plots on the bottom are empirical cumulative distribution function (CDF) plot highlights consensus distributions for each k.

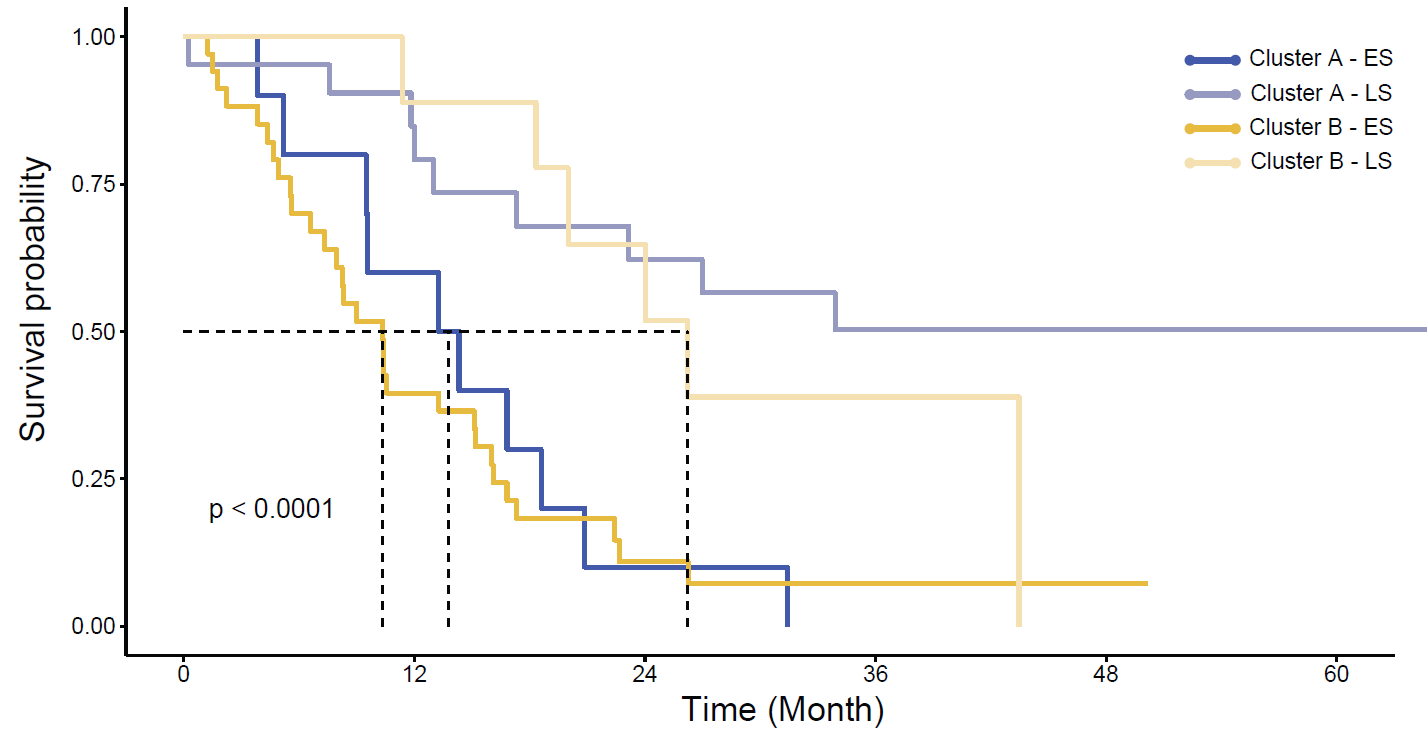

**Supplemental Figure 7.** Kaplan-Meier survival analysis on cluster A and B identified by consensus clustering stratified by either limited-stage or extensive-stage SCLC patients.

**Supplemental Table 1. Demographics by staging of SCLC patient cohort & non-cancer control participants. To test for association between (p values: continuous = Mann-Whitney & categorical = Fishers exact)**

| **Small cell lung cancer patients** | | | | | | **Non-cancer** |
| --- | --- | --- | --- | --- | --- | --- |
| **label** | **levels** | **Extensive-stage** | **Limited-stage** | **Total** | **p** |  |
| Total N (%) |  | N = 44 (59.5) | N = 30 (40.5) | N = 74 |  | N = 20 |
| Age (years) | Median (IQR) | 65.7 (60.8 to 74.5) | 68.8 (61.9 to 75.0) | 67.4 (61.3 to 75.1) | 0.60 | 68 (70.0 to 78.0) |
| Sex | Female | 12 (27.3) | 16 (53.3) | 28 (37.8) | 0.03 | 12 (60%) |
|  | Male | 32 (72.7) | 14 (46.7) | 46 (62.2) |  | 8 (40%) |
| Ethnicity | Asian | 7 (20.0) | 2 (6.9) | 9 (14.1) | 0.36 |  |
|  | Caucasian | 25 (71.4) | 25 (86.2) | 50 (78.1) |  |  |
|  | Other | 3 (8.6) | 2 (6.9) | 5 (7.8) |  |  |
|  | (Missing) | 9 | 1 | 10 |  |  |
| Smoking Status | Current smoker | 20 (45.5) | 13 (43.3) | 33 (44.6) | 0.39 | 20 (100%) |
|  | Former smoker | 21 (47.7) | 12 (40.0) | 33 (44.6) |  | - |
|  | Never smoker | 3 (6.8) | 5 (16.7) | 8 (10.8) |  | - |
| Smoking Pack Years | Median (IQR) | 50.0 (30.0 to 60.0) | 40.0 (30.0 to 50.0) | 40.0 (30.0 to 54.2) | 0.56 | 68 (59.7 to 91.7) |
| Thoracic Radiation | No | 15 (34.1) | 4 (13.3) | 19 (25.7) | 0.06 | - |
|  | Yes | 29 (65.9) | 26 (86.7) | 55 (74.3) |  |  |
| Any Systemic Therapy | No | 0 (0.0) | 2 (6.7) | 2 (2.7) | 0.16 | - |
|  | Yes | 44 (100.0) | 28 (93.3) | 72 (97.3) |  |  |

### **Supplemental Table 2. Two-variable Cox-proportional hazard model**

| Dependent: Survival (time_os_mo, event) |  | all | HR (univariable) | HR (multivariable) |
| --- | --- | --- | --- | --- |
| clusters | A | 31 (100.0) | - | - |
|  | B | 43 (100.0) | 2.02 (1.15-3.55, p=0.014) | 1.25 (0.68-2.29, p=0.470) |
